# Unexpected events modulate context signaling in VIP and excitatory cells of the visual cortex

**DOI:** 10.1101/2024.05.15.594366

**Authors:** Farzaneh Najafi, Simone Russo, Jérôme Lecoq

## Abstract

The visual cortex predicts incoming sensory stimuli through internal models of the world. Unexpected stimuli that violate these predictions update internal models and drive adaptation. Cortical inhibitory neurons, particularly VIP (vasoactive intestinal peptide) interneurons, are suggested to play a key role in representing unexpected stimuli, given their robust firing following unexpected omissions of familiar images. Importantly, this response is stimulus non-specific, raising an important question about what information it conveys. Given their unique connectivity with other cell types and brain areas, we hypothesized that during unexpected events, VIP neurons encode contextual information, defined as neuronal activity that is not driven by the stimulus itself. To test this hypothesis, we analyzed the Allen Institute Visual Behavior dataset, in which mice viewed repeated familiar images and unexpected omissions of these images while two-photon calcium imaging data from different cell types and visual areas were recorded. Using dimensionality reduction techniques, we found that omissions trigger contextual signaling in VIP neurons in the primary visual cortex (V1) and the lateral medial (LM) visual area, particularly in the superficial layers. Similarly, contextual coding was enhanced in excitatory neurons following omissions. This contrasted sharply with the excitatory response to expected images, during which contextual information was substantially suppressed. Our results suggest that unexpected events activate VIP neurons, which subsequently propagate contextual information across the cortical network. This potentially facilitates the integration of context within the cortical network, and leads to updated predictions about our dynamic environment.

## INTRODUCTION

The visual cortex is crucial in shaping our internal visual representations of the sensory environment. In the mouse visual cortex, inhibitory neurons are diverse, with major classes including parvalbumin (PV), vasoactive intestinal polypeptide (VIP), and somatostatin (SST) neurons, each with distinct subclasses as evidenced by transcriptomic characterization^1^. VIP cells, presumed to exert a dis-inhibitory effect on pyramidal cells, may play a key role in regulating information flow within the cortical network due to their connectivity with associative cortical areas and neuromodulatory systems^2–5^. VIP neurons exist in superficial (layer 2/3 primarily), and deeper (layer 4) layers of the cortex, suggesting their involvement in both top-down and bottom-up information processing, respectively^6^.

Circuit models suggest that unexpected stimuli trigger computations across both excitatory and inhibitory cortical cells to refine sensory representations^7^. Empirical data indicate that unexpected stimuli elicit responses across multiple layers of the cortex, involving inhibitory neurons^8^. A recent study demonstrated that the activity of VIP cells in the visual cortex peaks during unexpected omissions of visual stimuli^9^, but the underlying nature of these responses remains unclear. VIP cells may signal the presence of a prediction error signal; alternatively, they may encode contextual changes that co-occur during omissions, including behavioral, external, or internal states.

In the present study, we addressed this question using multi-area two-photon calcium imaging during a visually guided behavior. This task contained both expected events (regular flashes of images) and unexpected events (image omissions). We leveraged population activity decoding, dimensionality reduction and trial shuffling to examine whether VIP activity during omissions may contain stimulus-specific prediction-error signals, or if it was primarily influenced by context information. Importantly, “context” in our study broadly refers to shared activity across brain areas that are independent of task-relevant features. This may involve behavioral features such as the animal’s locomotion, external inputs such as the sensory signals in the animal’s environment, and internal representations related to the animal’s current state.

We found that most VIP neurons, except those in layer 4 of the primary visual area, encoded context information. This suggests that VIP neurons represent a general ‘surprise’ signal enriched with contextual details during unexpected events. In contrast, excitatory neurons displayed weaker modulation by context during expected image presentations but were significantly influenced by context during unexpected omissions. These results highlight the role of superficial VIP neurons in updating cortical activity with contextual information in response to unexpected stimuli. This mechanism potentially facilitates the association of contextual signals with unexpected events, thereby driving adaptive responses.

## RESULTS

Distinct cortical pathways are thought to carry sensory and prediction signals^10,11^, suggesting that cortical areas and cell types might show unique responses to expected vs. unexpected events. To test this hypothesis, we leveraged two-photon imaging data from the Allen Brain Observatory platform^9,12^. In this dataset, mice were trained to perform a go/no-go, image-change detection task^9^. In brief, water-restricted animals were presented with a constant stream of natural images (250 ms) interleaved with gray screens of matched luminance (500 ms). When a change in image identity occurred, mice would receive a water reward if they licked immediately after the image change. Importantly, during imaging sessions in well-trained mice, a pseudo-random 5% of non-change images were replaced with a gray screen (image omissions; Fig. 1A). Since mice had extensive experience with the highly regular timing of image presentations (every ∼750 ms), the rare stimulus omissions were unexpected events, as opposed to the expected, frequent, and repeated images. During this task, the activity of excitatory neurons as well as inhibitory SST and VIP neurons were recorded separately, using 3 distinct mouse lines (see Methods). Neuronal cells within each line were simultaneously imaged at 4 cortical depths in 2 visual areas (V1 and higher visual area: LM; Fig. 1B,C; Supplementary Fig. 1).

**Figure 1.**
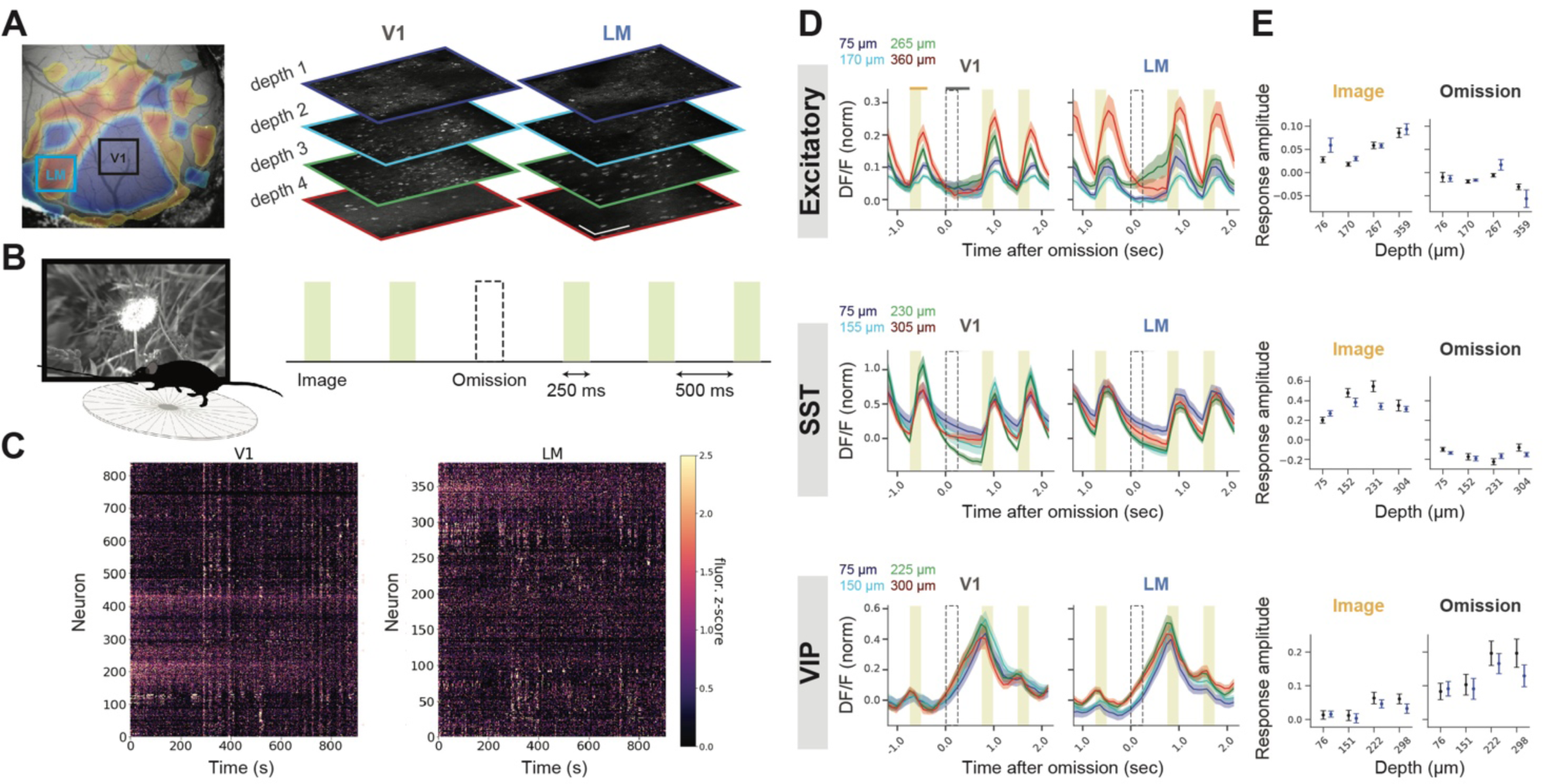
Excitatory and inhibitory cell types distinctly encode expected images and unexpected omissions across visual areas and layers. **A)** Allen Institute Visual Behavior dataset: calcium signals were simultaneously imaged across two cortical columns, including 4 depths (approximately 100 µm, 200 µm, 300 µm, and 375 µm), in visual areas V1 and LM, and in 3 distinct mouse lines tagged for excitatory, and 2 inhibitory cell types: VIP and SST. B) Mice, walking on a disk, view a familiar set of images, separated by a gray screen. Images are unexpectedly omitted on 5% of the presentation. C) Example population activity of excitatory neurons, recorded simultaneously from ∼1180 neurons in V1 and LM. D) Population-averaged calcium responses to images (shaded rectangle) and omissions (dashed rectangle), across 2 visual areas (V1, LM), and 4 cortical depths (blue, cyan, green, red), in 3 different mouse lines (A-C: excitatory, SST, VIP). ΔF/F traces normalized to baseline standard deviation. Mean +/- SEM; n = 24, 22, 24 sessions for excitatory, SST, and VIP, respectively. E) Quantification of evoked responses averaged over 350ms after images, and 500ms after omissions (time-windows used for quantification of image and omission-evoked responses are indicated by orange and gray horizontal lines above top left panel).

We first studied how excitatory and inhibitory subtypes (VIP, SST) respond to image presentations and omissions, and how their responses depend on cortical area and depth. The population average of neural activities demonstrated a clear difference across cell classes (Fig. 1D,E). Excitatory neurons and, more robustly, SST neurons in all areas and layers were activated by expected images, i.e. the expected events. Conversely, excitatory neurons, overall, did not respond to omissions, except for those in the middle depth of LM. SST neuronal activity was slightly decreased after omissions in all recorded locations (Fig. 1D,E top, middle). In sharp contrast to excitatory and SST neurons, VIP neurons in all layers and areas were robustly activated after omissions (Fig. 1D,E bottom). Specifically, VIP cells demonstrated small ramping activity approximately 250 ms prior to each image presentation that further increased when the image was not presented (i.e. omission), and was immediately inhibited by image presentation, confirming previous results using the same behavioral task^9^.

This multi-area dataset allowed comparison of neuronal responses across cortical depths of V1 and LM in each experiment (Fig. 1D,E). We found that excitatory neuronal responses to images were stronger in deeper layers of both visual areas, particularly V1. SST responses to images were strongest at middle depths of V1, corresponding to layer 2/3 and superficial layer 4, but did not differ among cortical layers of LM. VIP responses to neither images nor omissions were significantly different across cortical layers or areas, as reported previously^6^, although the responses were slightly stronger in deeper layers. Therefore, in both V1 and LM, images are more robustly represented by excitatory and SST neurons, particularly in deeper layers; while unexpected omissions are represented by VIP neurons (Fig. 1D,E; Stats: Supplementary Table 1).

It is suggested that feedback pathways, from higher order cortical areas to superficial layers of V1, convey prediction signals^10,11,13^. In light of these studies, we used simultaneous multi-plane recording to investigate if cortical interactions might be different when expectations are violated, i.e. during omissions. As a proxy for cortical interactions, we measured pairwise noise correlations across repeated image presentations, as well as omissions, for each cell type. Correlations were computed within each area (V1-V1, LM-LM; Supplementary Fig. 2-4), as well as across areas (V1-LM; Fig. 2). We also performed a shuffling analysis to control for firing rate impacts on correlations (Methods; Supplementary Fig. 2-4)

**Figure 2.**
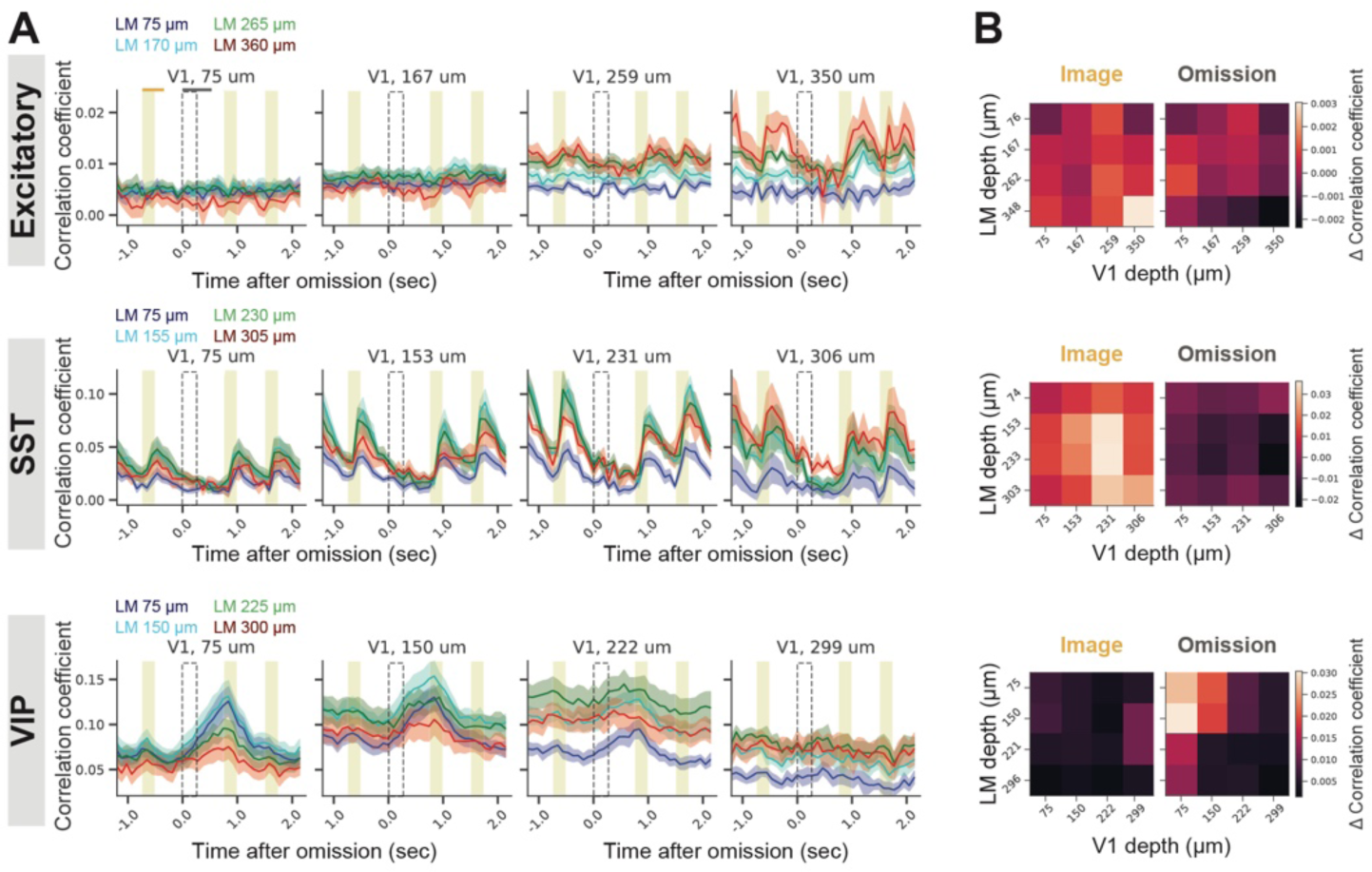
Distinct cortical interactions occur across primary and higher-order visual areas following expected and unexpected events. **A)** Spearman correlation coefficients between V1 and LM neurons at different cortical depths (colors indicate LM depths; each subplot corresponds to a given V1 depth). **B)** Change in correlation coefficients during images (left) and omissions (right) relative to baseline correlation coefficient. Correlation coefficients quantified over 500ms after images, and 750ms after omissions. Traces and heatmaps: mean +/- SEM; n = 8, 6, 9 mice for excitatory, SST, and VIP cell types, respectively.

In the excitatory network, V1-V1 and V1-LM coactivations increased after image presentations, but only in deep layers (V1-LM: Fig. 2; V1-V1: Supplementary Fig. 2A,D). Interestingly, excitatory network coactivation in LM-LM did not change after images, across any pair of cortical layers (LM-LM: Supplementary Fig. 2B,E). Following omissions, there was no significant change in the coactivation of excitatory neurons across visual layers and areas (Fig. 2; Supplementary Fig. 2A,D).

In the SST network, images increased coactivation more broadly across visual layers and areas, particularly in our “middle” imaging depth, i.e., deep layer 2/3 and superficial layer 4 (Fig. 2; Supplementary Fig. 3). Omissions, on the other hand, slightly decreased coactivation of neurons relative to the baseline across most layers of the SST network (Fig. 2; Supplementary Fig. 3).

In the VIP network, coactivation patterns were strikingly different compared to other cell types: neurons in *superficial* layers of V1 and LM were strongly coactivated following omissions. There was also a weak coactivation prior to images in most layers and areas (Fig. 2; Supplementary Fig. 4). Notably, correlations were overall much weaker among excitatory neurons compared to inhibitory neurons (Fig. 2), confirming previous results and indicating stronger local connectivity between inhibitory neurons^14,15^.

Our results suggest that the VIP network in the visual cortex is strongly recruited during unexpected events: cells from all layers of V1 and LM are active during omissions and neural correlations are especially strong in the superficial layers. One possible explanation is that prediction errors (i.e. mismatch between the animal’s expectation of a given image and its omission) generate a shared input signal that co-activates VIP cells in superficial layers across areas, thereby synchronizing the local cortical network. An alternative hypothesis is that VIP activity during omissions may be due to co-occurring changes triggered by unexpected events. Such changes could include variations in animal’s locomotion and arousal, as well as alterations in the representation of environmental cues, or internal states. This hypothesis is corroborated by recent reports^16,17^, including our own^18^, that have demonstrated a broad range of behavioral features modulate brain-wide network dynamics, raising the possibility that our correlation findings could be contributed by global brain fluctuations driven by widespread behavioral events. Importantly, in our previous work ^9^, we had found that the VIP population carries no information about the identity of the previous image. These results demonstrated that VIP neurons do *not* encode stimulus-specific prediction error signals; instead suggesting that VIP omission activity may represent a broad non-specific surprise signal.

To address whether VIP correlated activity during omissions may be affected by global contextual changes, we performed Tensor Component Analysis (TCA)^19^ on real and trial-shuffled data, aligned on images and omissions (number of sessions = 22, 15, 21 for excitatory, SST, and VIP cells, respectively). TCA allowed decomposing neural population activity into distinct components, and trial shuffling allowed studying contextual representation (see below). Each TCA component was characterized by weights indicating the unique contributions from individual neurons, trials, and time points (Fig. 3A,B). Importantly, we defined TCA components as image-specific or omission-specific if their trial weights were significantly higher on images or omissions, respectively (ranksum test across image and omission weights; p < 0.05; Fig. 3E). Components in which trial weights were not significantly different between images and omissions were excluded from further analyses (number of non-specific components, pooled across sessions; mean +/- SD across 100 TCA iterations; Excitatory: 38 +/- 4; SST: 20 +/- 3; VIP: 55 +/- 3).

**Figure 3.**
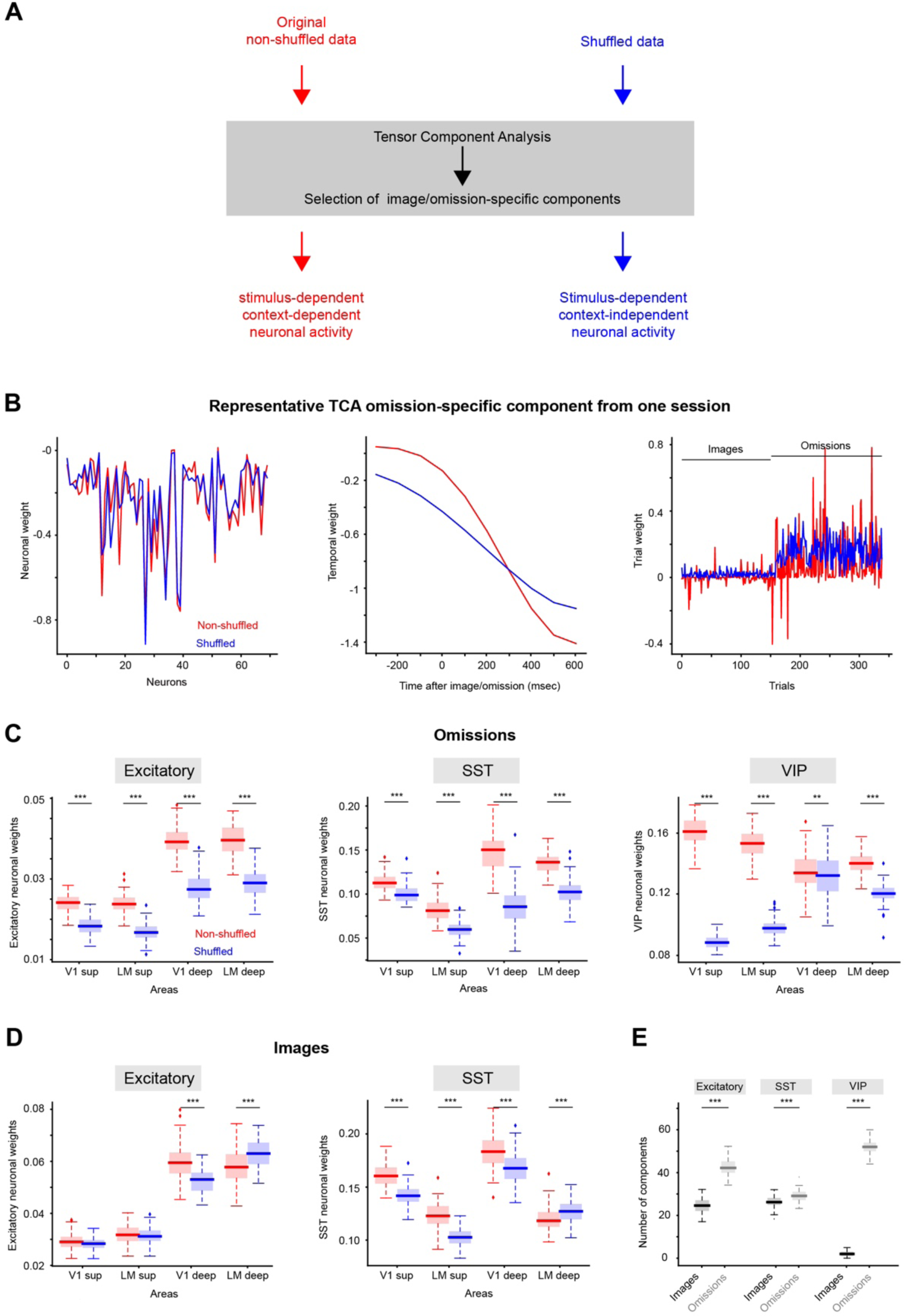
Unexpected events lead to enhanced context representation in excitatory and particularly VIP neurons in superficial layers. **A)** Schematic of the analysis pipeline. Imaging data were decomposed using Tensor Component Analysis (TCA; red: original; blue: shuffled). Shuffled data were obtained by permuting trials identities within each category (image and omission). Each TCA component was identified as image-specific, omission-specific, or non-specific depending on the weight magnitude of the TCA trial component for images vs. omissions (Methods). **B)** Left to right: neuronal, temporal, and trials weights of a representative omission-specific TCA component (red: original; blue: shuffled). In this example, trials from 1 to 159 are images, while the remaining trials are omissions. TCA was performed on spikes extracted from DF/F traces aligned on images and omissions, including 300ms before to 700ms after images and omissions. **C)** Neuronal weight of the omission-specific components for excitatory, SST, and VIP neurons (left to right) extracted from original and shuffled data across areas (Wilcoxon ranksum test; all p<0.001 except for VIP V1 deep: p=0.009). **D)** Same as panel C for image-specific components for excitatory and SST neurons (left to right; Wilcoxon ranksum test; all p<0.001 except for excitatory V1 superficial: p=0.8 and excitatory LM superficial: p=0.5). **D)** Number of components pooled across sessions, for excitatory, SST, and VIP neurons (left to right) from original data compared between images and omissions (black and gray boxes, respectively; Wilcoxon ranksum test; all p<0.001).

To investigate the neural coding of images and omissions across visual areas and layers, we compared the absolute magnitude of TCA neuronal weights among V1, LM, superficial and deep layers (imaging depth smaller or bigger than 250μm, respectively; Fig 3C,D, red; for the details of TCA stats see Table S2). In omission-specific components, VIP neurons had stronger weights in *superficial* layers of both V1 and LM. This contrasted with excitatory and SST neurons which had stronger weights in deeper layers of V1 and LM (Fig. 3C, red). In image-specific components, excitatory neurons in deeper layers of V1 and LM had stronger weights; while SST neurons showed stronger weights in V1 compared to LM, independent of cortical layer (Fig. 3D red). VIP neurons had few image-specific components, hence excluded from our analysis of neuronal weights.

TCA neural weights for image- and omission-specific components reflect the coding strength of individual neurons on image and omission trials, respectively. This neural coding could arise from the direct presentation of images and omissions, or could be influenced by co-occurring behavioral and sensory features in adjacent or remote areas, which we collectively term contextual features. To distinguish the neuronal coding of images and omissions from these contextual influences, we performed TCA on trial shuffled data, for which trial identities were shuffled *within* the image and omission conditions for each neuron (see Methods). This shuffling procedure allowed us to preserve image- and omission-specific responses while removing any contextual information that co-occurred across neurons during images and omissions. In contrast, in the non-shuffle data, TCA components could encode the presence of images and omissions as well as any context-dependent information present in the neuronal population.

Shuffling led to a general decrease of neuronal weight magnitudes for omission-specific components in all cell types (Fig. 3C blue: shuffled; red: real data). This reduction was most prominent in VIP neurons in the superficial layers, and absent from the VIP neurons in deep V1 (Fig. 3C, right). These findings suggest that omissions are co-represented with contextual information in the visual cortex in excitatory, SST, and particularly superficial VIP neurons. The major exception is that VIP neurons in the deeper layers of V1 show a smaller contribution from contextual signals (Wicoxon ranksum test; p=0.009).

Interestingly, shuffling had a smaller effect on the neuronal weights of image-specific components in excitatory and SST cells (Fig. 3D blue vs. red; Note the absence of VIP neurons from this analysis due to their few numbers of image components). This finding suggests that during images, excitatory neurons primarily encode image information, overriding contextual signals. This is aligned with previous studies that demonstrated reduced neuronal variability during sensory inputs^20^.

Indeed, all cell types, and particularly VIP neurons, had more TCA components that were omission-specific vs. image-specific (Fig. 3E, black: images, gray: omissions). This finding suggests that omission responses in our neuronal population exhibit more variability than image responses and is aligned with our TCA shuffled analysis which demonstrated response variability during omissions was associated with the co-representation of contextual features.

## DISCUSSION

Recent work had demonstrated that the VIP population is functionally quite diverse, and strongly represents behavioral features and unexpected image omissions^9^. Extending these findings, our current study demonstrates that during unexpected omissions, VIP neurons in the visual cortex not only encode a non-specific surprise signal but also a substantial amount of contextual information. This dual representation of omissions and context is predominantly observed in superficial VIP neurons, with notably reduced context coding in deeper layers.

Multiple lines of work have suggested the role of VIP neurons in encoding external and internal features. VIP neurons can represent behavioral state, leading to gain modulation in sensory cortices^4,21^. VIP activity increases with pupil dilation^22^, locomotion^4,23^, and whisking ^3,24^, leading to enhanced sensory modulation during high attention and arousal states that follow surprising and reinforcement signals^4,21–23,25,26^. VIP neurons are driven by neuromodulators, particularly cholinergic inputs^4^, as well as long-range cortical and thalamic inputs^2,3^. VIP neurons have also been shown to play a key role to surround suppression^27^, when neuronal responses to centered visual stimuli can be decreased by the presence of adjacent stimuli.

Our study suggests that the VIP network in primary and higher visual cortices comprises two subnetworks with distinct roles, associated with superficial and deeper layers. In V1, the subnetwork in the deeper layers encodes information about omissions, independently of context, while the subnetwork in the superficial layers concurrently encodes both context and omission information. Importantly, the omission coding by VIP cells is non-specific, lacking any information about which stimulus was omitted, as suggested by our recent work (Garrett 2023). This supports the hypothesis that VIP neurons broadly encode surprise signals rather than specific mismatches between expected and actual stimuli. Our findings suggest a potential specialization within the VIP network that enhances the reliable representation of sensory features in deeper V1, while integrating sensory and contextual features in the superficial layers of V1, hence facilitating adaptive responses to dynamic environmental stimuli.

The ability of VIP cells to disinhibit excitatory neurons through its connectivity pattern, is ideally suited to facilitate the propagation of selective activity patterns through the cortex, leading to attention towards contextual sensory inputs^26^ when needed, or the emergence of motor behaviors^28,29^. We propose a framework in which VIP cells serve as gatekeepers of contextual information, which are dynamically activated when transitioning from expected to unexpected stimuli (Fig. 4). This framework is in agreement with the numerous studies on the activity of the inhibitory network. Future experiments should aim to simultaneously record from all those cell types in the context of dynamic but controlled sensory uncertainty.

**Figure 4.**
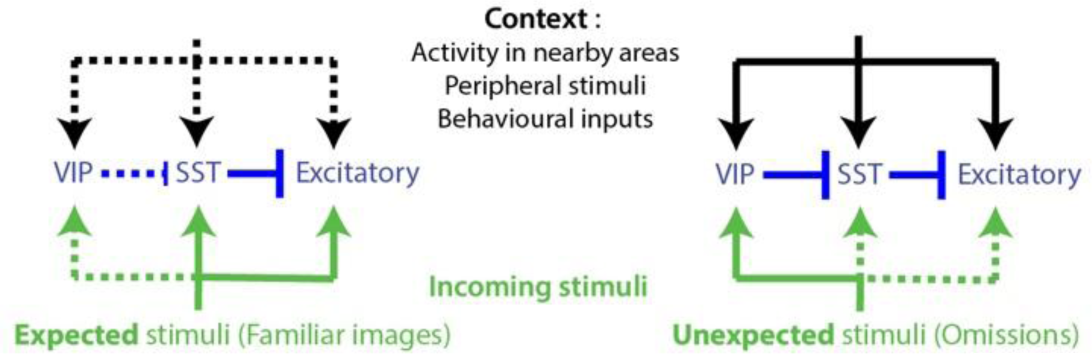
Proposed mechanism: Triggered by unexpected events, VIP neurons gate contextual signal flow across cortical cell types and areas. **Left)** Expected stimuli, such as familiar images in our study, are encoded by excitatory and SST neurons. **Right)** When unexpected stimuli occur, a non-specific surprise signal activates VIP cells to promote contextual integration across all cell types. This contextual information may originate from nearby and distant brain areas, encoding a variety of signals, including internal (cognitive and motivational states), or external (sensory and behavioral) information. The presence of this contextual information facilitates the updating of the unexpected stimulus representation in the local network.

## ONLINE METHODS

### Processing of calcium imaging movies

The preprocessing of all calcium imaging data was done within the imaging processing pipeline described in detail elsewhere^34^ and crosstalk removal was applied after pre-processing. Briefly, all data was first corrected for brain motion by performing rigid registration in two dimensions. Then, cell-segmentation was performed to identify spatial masks of active neurons. Further, fluorescence from spatially-overlapping neuronal masks was unmixed and corrected for neuropil contamination. Finally, mask-matching and crosstalk removal was performed using FastICA^37^ and ghost cells were filtered, and lastly, ΔF/F was computed on corrected masks.

### Description of datasets

We used the publicly available Allen Institute Visual behavior dataset: https://portal.brain-map.org/circuits-behavior/visual-behavior-2p. Details are explained in our previous work (Garrett 2023). In brief, this dataset included 3 mouse lines: excitatory mouse line: Slc17a7-IRES2-Cre; Camk2a-tTA; Ai93(TITL-GCaMP6f), inhibitory subpopulation VIP: Vip-IRES-Cre; Ai148 (TIT2L-GC6f-ICL-tTA2), and inhibitory subpopulation SST: Sst-IRES-Cre; Ai148 (TIT2L-GC6f-ICL-tTA2). For the excitatory cell line, 24 sessions from 8 mice; for the SST cell line, 22 sessions from 6 mice; for the VIP cell line, 24 sessions from 9 mice were recorded.

### Population averages of neuronal responses

Traces were aligned to the onset of image omission; since images were presented regularly, omission alignment also aligned the traces on images. The median of neural responses was computed across trials. The average response was then computed across neurons, for each session. A grand average was then computed across sessions (see Fig. 1D, traces). To quantify the image-evoked responses, the calcium trace of each neuron was averaged over 350ms after image onset for excitatory and SST neurons, and over [-250, 100]ms relative to image onset for VIP neurons to account for their anticipatory response. Response quantification was done on the mean (across trials) trace of each neuron. The same quantification was performed for omission-evoked responses, except a 500ms window was used for quantification. The responses were averaged across neurons, for each session, and then a grand average was computed across sessions (see Fig. 1E; error-bars).

### Correlation of neural responses across cortical planes

On omission-aligned traces, Spearman correlation coefficient was computed between pairs of neurons across trials. Here, by trial we mean omissions. Correlations were computed for every individual frame, over [-1, 2] sec relative to the omission. This procedure was done for all pairs of neurons; then an average value was computed across all pairs (Fig. 2). Neuron pairs were present within the same plane, or in 2 different planes. This analysis allowed studying how the response of neurons (within the same plane or in different areas/layers) covaried across trials, and how this coactivation changed at different moments (e.g. after images vs. omissions). Spearman correlation coefficients were also computed on shuffled traces (Supplementary Fig. 2-4 d-f), which were driven by independently shuffling trial orders for each neuron. For each neuron pair, shuffling was repeated 50 times, resulting in a distribution of correlation coefficients for shuffled data. This shuffling analysis controlled for firing rate impacts on our correlation results.

<u>T</u>o compute “noise” correlations, we measured pairwise correlations of “signal”-removed traces. “Signal” was computed by taking the average neural response to each image type (there were 8 images in each session), and subtracting the average from the response of individual trials of that image type.

To quantify coactivation of neurons (Fig. 2, heatmaps; Supplementary Fig. 2-4d-f), correlations coefficients (“cc”) were first averaged over 500 ms after images for excitatory and SST neurons, and over [-250, 250] ms relative to image onset for VIP neurons, accounting for their anticipatory response. Then, correlation coefficients were averaged across baseline frames. We call this quantity “image cc”. To quantify omission-evoked coactivation, we averaged correlation coefficients over 750 ms after omissions. We call this quantity “omission cc”. Next, we quantified baseline coactivation by averaging correlation coefficients across baselines frames. Baseline was defined as the frame immediately preceding each image presentation, for excitatory and SST neurons, and 250 ms earlier than each image presentation for VIP neurons. We call baseline quantification of correlation “baseline cc”. Finally, we measured the change in coactivation during images or omissions by subtracting out “baseline cc” from “image cc”, or “omission cc”. This quantity is plotted in Fig. 2, heatmaps, and Supplementary Fig. 2-4d-f, error-bars.

The Python package scipy (scipy.stats.spearmanr) was used for computing correlations, and p-values were computed in 2 ways: 1) using the p-value output of the spearmanr package; 2) manually computing the p-value by comparing the correlation coefficient of real (non-shuffled) data with the shuffled distribution using 2-sided, 1-sample t-test (using the Python package scipy.stats.ttest_1samp).

### Tensor Component Analysis

Tensor Component Analysis (TCA) was performed on inferred spikes from the DF/F trace using the tensortool package (https://github.com/neurostatslab/tensortools). TCA was set to extract 5 components from the trials split from -300 to +600 msec relative to the image or omission onset. The resulting components were first filtered according to their image- or omission-specificity. Specifically, we compared the weights of image and omission trials for each component (Wilcoxon ranksum test, alpha <0.05) and identified as condition-specific the components showing a significant difference between the two. On this subset of components, those with larger absolute average weights during image trials were classified as image-specific, while those with larger absolute average weights during omission trials were classified as omission-specific. To assess the variability across trials, on each subset of components we computed the standard deviation across trial weights. This entire procedure was performed 100 times to account for the variability between TCA decompositions. To obtain a null distribution that discarded context information while preserving image- and omission-specific information, we shuffled trials for each neuron keeping separated image and omission trials (e.g. the activity of an omission trial was reassigned to another omission trial independently for each neuron). Similarly to the original data, the procedure was performed 100 times to account for variability in shuffling and TCA decomposition. While TCA explained a relatively small amount of the overall signal, especially in the case of excitatory neurons, the components were highly reproducible across iterations (for TCA error and similarity across iterations see Table S3). As an additional control, we performed the same analysis with a larger number of reconstructed components by TCA and found similar results (Supplementary Fig. 5).

### Statistical tests

We used two-way ANOVA, followed by Tukey HSD to compare population averages across cortical layers. Two-sided t-test was used to compare correlations between real and shuffled data, for each cortical plane. A p-value of 0.05 was used as the significance threshold. For comparison of the correlation coefficients, we used two-tailed, two-sample Kolmogorov–Smirnov tests.

## Data availability

Data are publicly available as Allen Institute Visual Behavior Dataset: https://portal.brain-map.org/circuits-behavior/visual-behavior-2p

## Code availability

Crosstalk removal was performed using custom routines employing FastICA which is available as part of scikit-learn python package (https://scikit-learn.org/). Code to perform population average and correlation analyses are available at: https://github.com/farznaj/multiscope_fn. Code to perform TCA and the related statistical tests are available at: https://github.com/SimoneRusso/Multiscope_TCA

## Acknowledgements

We wish to thank the founder of the Allen Institute for Brain Science, Paul G. Allen, for his vision, encouragement and support.

## Competing interests

The dual-beam add-on module (J.L.) intellectual property has been licensed to Thorlabs. Inc., by the Allen Institute. S.R. is the Chief Medical Officer of Manava Plus.

**Correspondence and requests** for materials should be addressed to J.L, and F.N..

## Supplementary material

**Supplementary Table 1.**
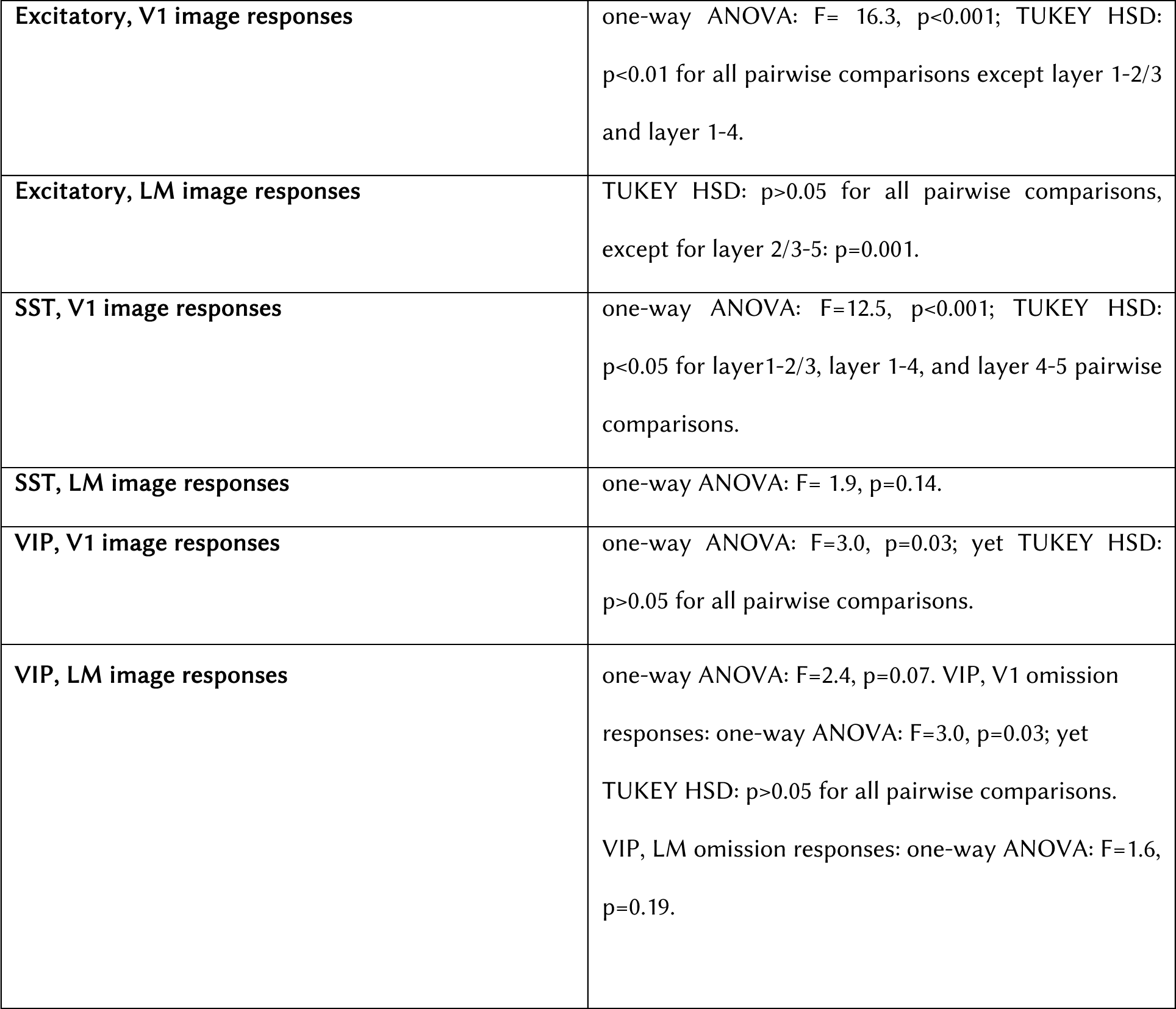
Statistical tests corresponding to Fig. 1.

|  |  |
| --- | --- |
| **Excitatory, V1 image responses** | one-way ANOVA: $F=16.3$, $p<0.001$; TUKEY HSD: $p<0.01$ for all pairwise comparisons except layer 1-2/3 and layer 1-4. |
| **Excitatory, LM image responses** | TUKEY HSD: $p>0.05$ for all pairwise comparisons, except for layer 2/3-5: $p=0.001$. |
| **SST, V1 image responses** | one-way ANOVA: $F=12.5$, $p<0.001$; TUKEY HSD: $p<0.05$ for layer 1-2/3, layer 1-4, and layer 4-5 pairwise comparisons. |
| **SST, LM image responses** | one-way ANOVA: $F=1.9$, $p=0.14$. |
| **VIP, V1 image responses** | one-way ANOVA: $F=3.0$, $p=0.03$; yet TUKEY HSD: $p>0.05$ for all pairwise comparisons. |
| **VIP, LM image responses** | one-way ANOVA: $F=2.4$, $p=0.07$. VIP, V1 omission responses: one-way ANOVA: $F=3.0$, $p=0.03$; yet TUKEY HSD: $p>0.05$ for all pairwise comparisons. VIP, LM omission responses: one-way ANOVA: $F=1.6$, $p=0.19$. |

**Supplementary Table 2.**
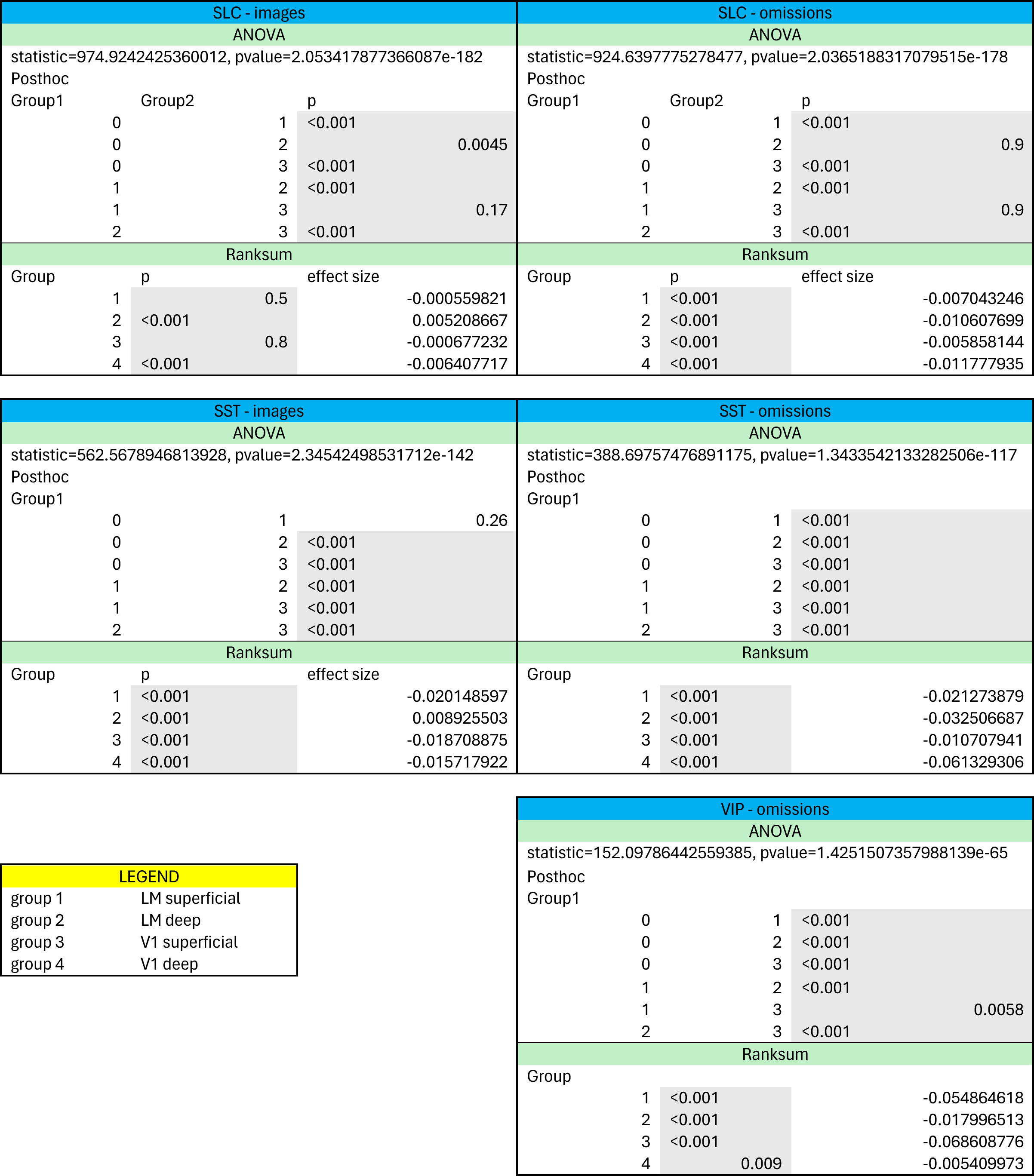
Statistical tables for TCA analysis.

**Supplementary Table 3.** TCA error and similarity values across with across number of components, celltypes, and original vs shuffled data.

| Number of components | Celltype | Data | Error | Similarity |
| --- | --- | --- | --- | --- |
| 5 | Excitatory | Original | 0.9401 | 0.8879 |
|  |  | Shuffled | 0.9681 | 0.8673 |
|  | SST | Original | 0.7695 | 0.9227 |
|  |  | Shuffled | 0.8365 | 0.9067 |
|  | VIP | Original | 0.774 | 0.918 |
|  |  | Shuffled | 0.8838 | 0.8886 |
| 10 | Excitatory | Original | 0.9084 | 0.8265 |
|  |  | Shuffled | 0.9366 | 0.8038 |
|  | SST | Original | 0.67 | 0.8695 |
|  |  | Shuffled | 0.7481 | 0.858 |
|  | VIP | Original | 0.6991 | 0.8515 |
|  |  | Shuffled | 0.8238 | 0.8195 |

**Supplementary Figure 1.**
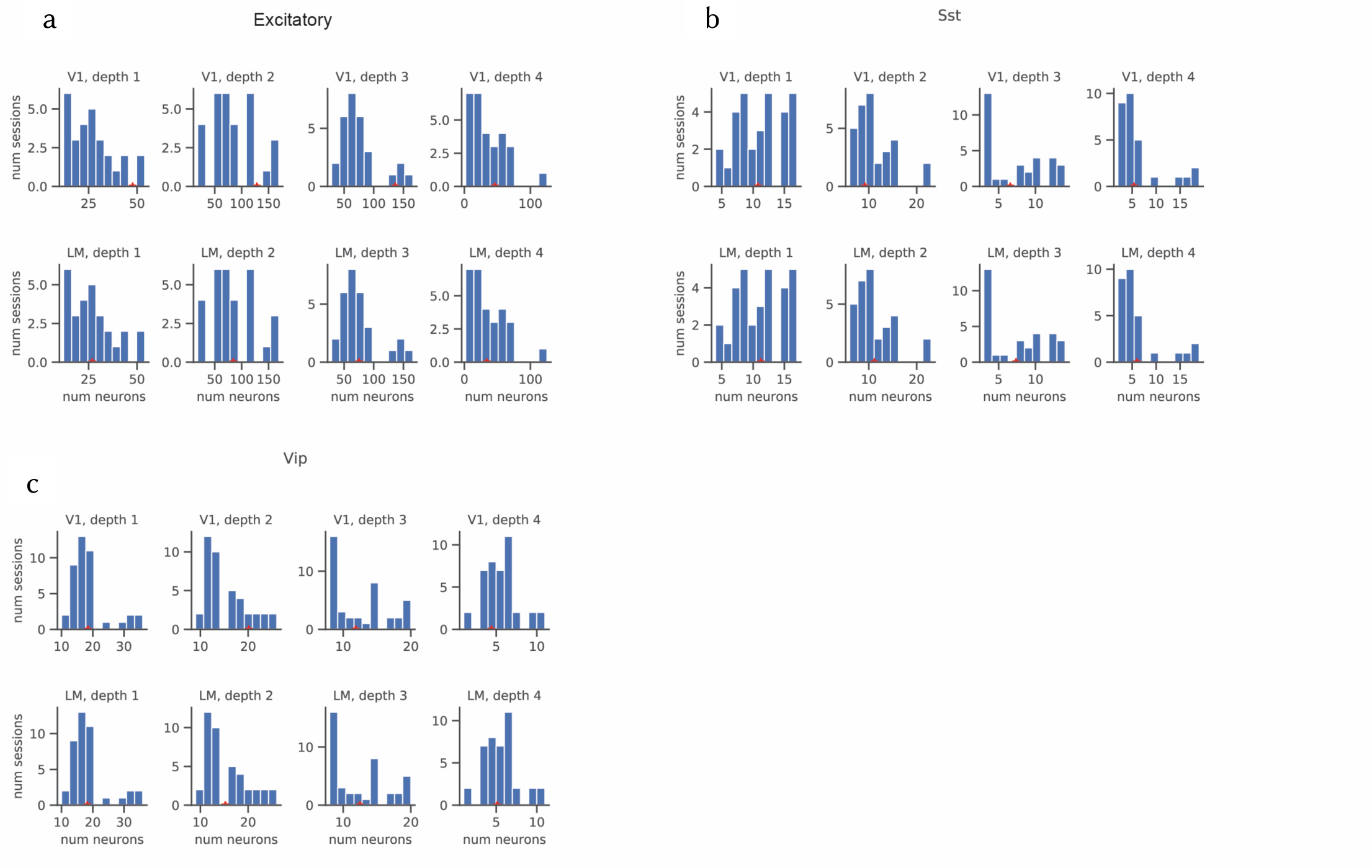
Number of neurons per depth and area, for each cell type. Distribution of number of neurons recorded simultaneously across depths of V1 (top) and LM (bottom), in all recording sessions, for excitatory **(a)**, SST **(b)**, and VIP **(c)** mice. Note that our goal in this study was to chronically record the activity of distinct excitatory and inhibitory cell types, from the same field of view and across multiple sessions. Like in other studies involving the original Mesoscope^1,2^, our system can record from a much larger number of neurons, however, our biological experiments were not designed for that purpose.

**Supplementary Figure 2.**
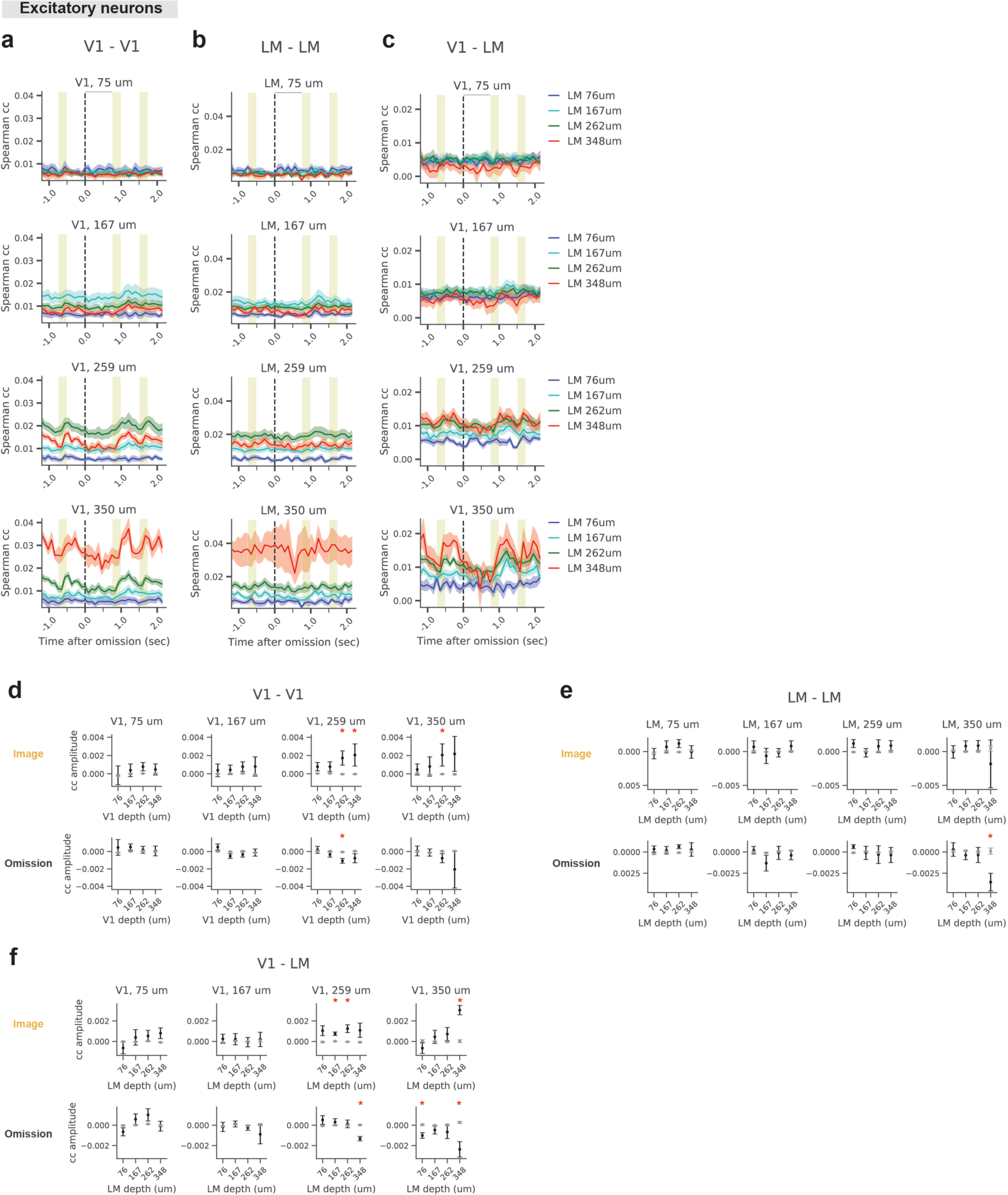
Correlation of excitatory neurons within V1, within LM, and across V1-LM, during images and omissions. **a-c** Spearman correlation coefficients computed, at different moments in the trial, between activities of excitatory neurons located in different depths of V1-V1 (a), LM-LM (b), and V1-LM (c). **d-f** Change in correlation coefficients during images (top) and omissions (bottom) relative to the baseline correlation coefficient, computed for the real data (black), and trial shuffled data (gray). Correlation coefficients were quantified over 500 ms after images, and 750 ms after omissions. Red stars indicate statistical significance (two-sided t-test, real vs. shuffle data; p<0.05). Traces and error-bars: mean +/- SEM; n = 8 mice.

**Supplementary Figure 3.**
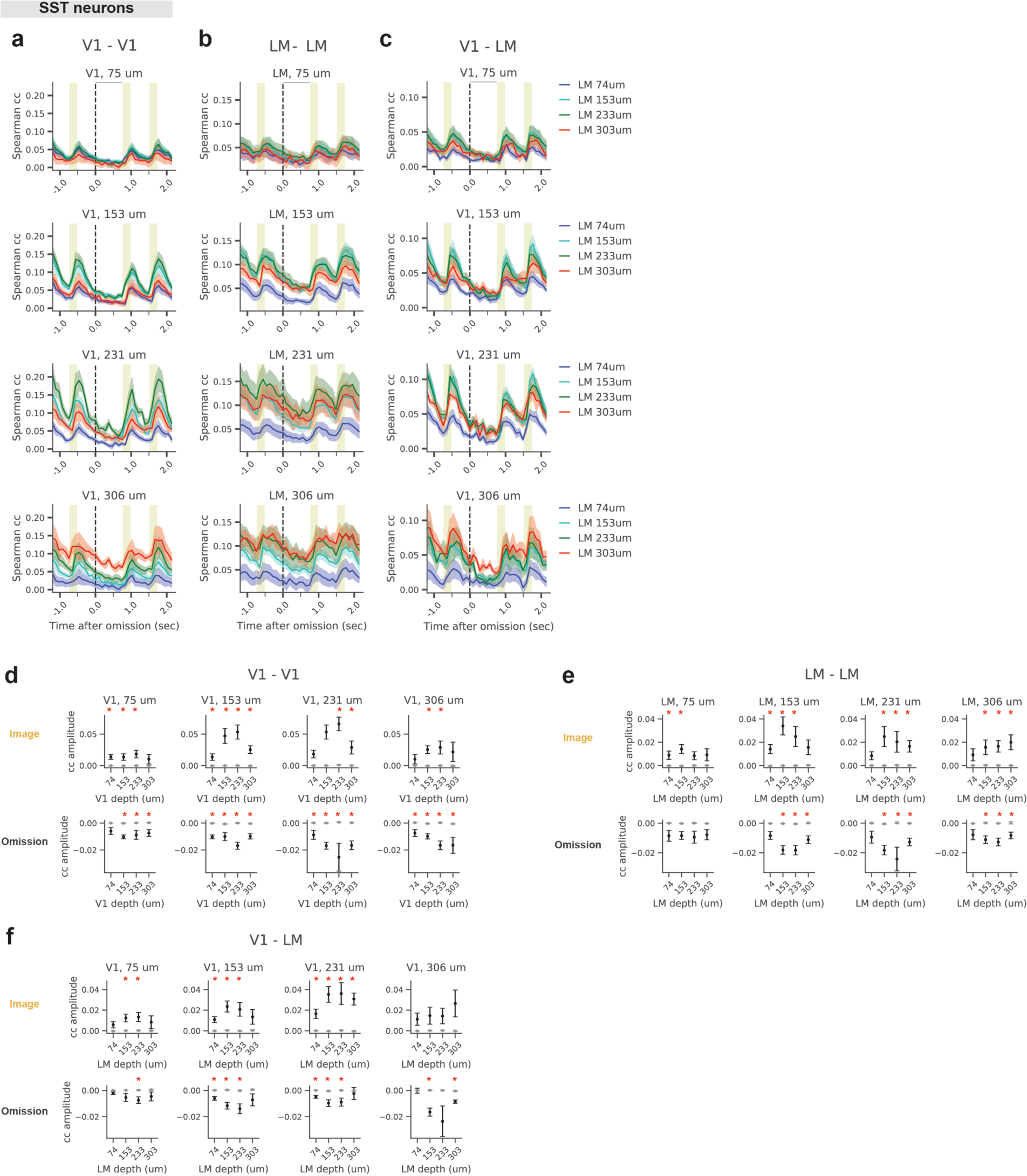
Correlation of SST neurons within V1, within LM, and across V1-LM, during images and omissions. **a-c** Spearman correlation coefficients computed, at different moments in the trial, between activities of excitatory neurons located in different depths of V1-V1 (a), LM-LM (b), and V1-LM (c). **d-f** Change in correlation coefficients during images (top) and omissions (bottom) relative to the baseline correlation coefficient, computed for the real data (black), and trial shuffled data (gray). Correlation coefficients were quantified over 500 ms after images, and 750 ms after omissions. Red stars indicate statistical significance (two-sided t-test, real vs. shuffle data; p<0.05). Traces and error-bars: mean +/- SEM; n = 6 mice.

**Supplementary Figure 4.**
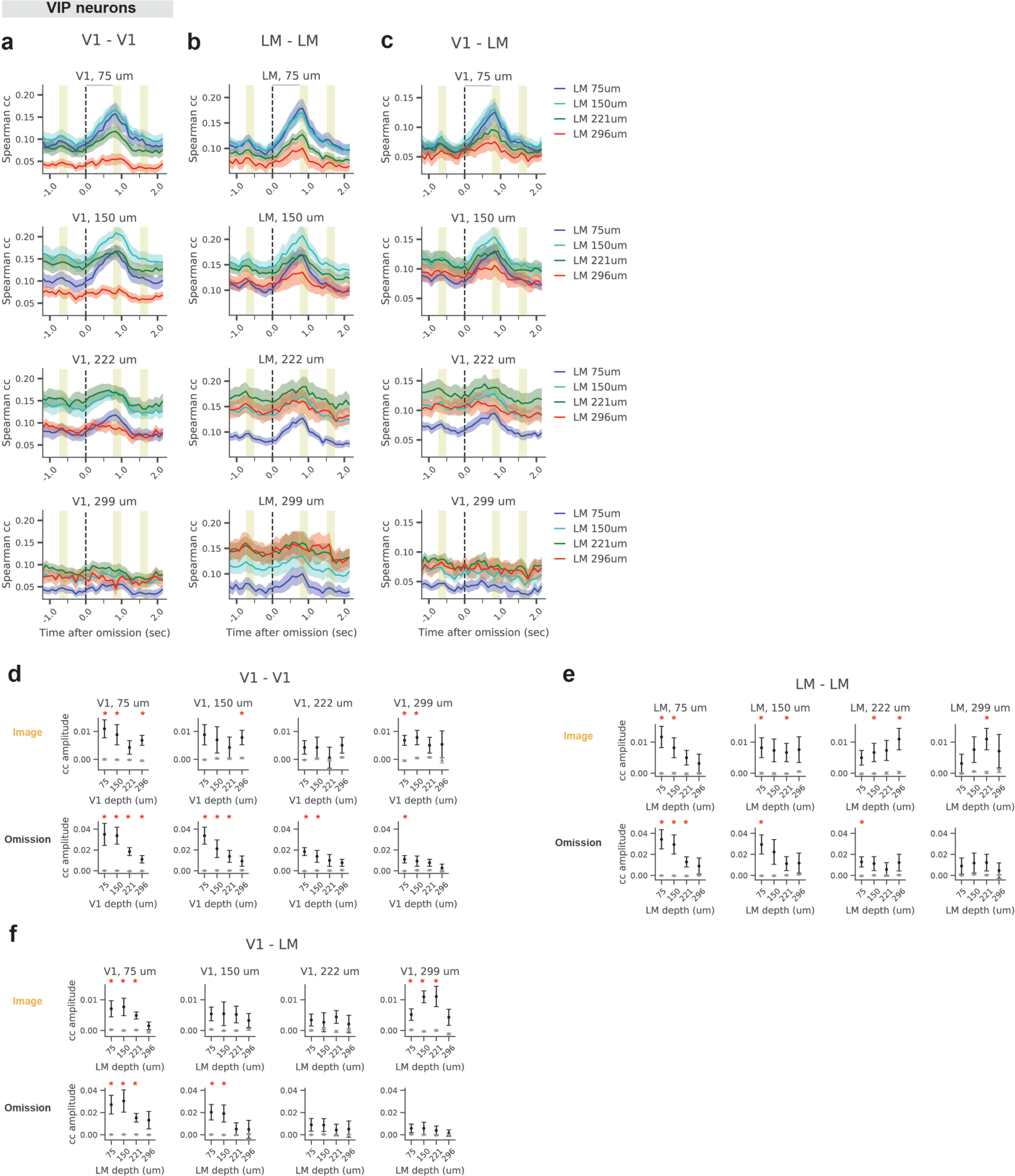
Correlation of VIP neurons within V1, within LM, and across V1-LM, during images and omissions. **a-c** Spearman correlation coefficients computed, at different moments in the trial, between activities of excitatory neurons located in different depths of V1-V1 (a), LM-LM (b), and V1-LM (c). **d-f** Change in correlation coefficients during images (top) and omissions (bottom) relative to the baseline correlation coefficient, computed for the real data (black), and trial shuffled data (gray). Correlation coefficients were quantified over 500 ms after images, and 750 ms after omissions. Red stars indicate statistical significance (two-sided t-test, real vs. shuffle data; p<0.05). Traces and error-bars: mean +/- SEM; n = 9 mice.

**Supplementary Figure 5.**
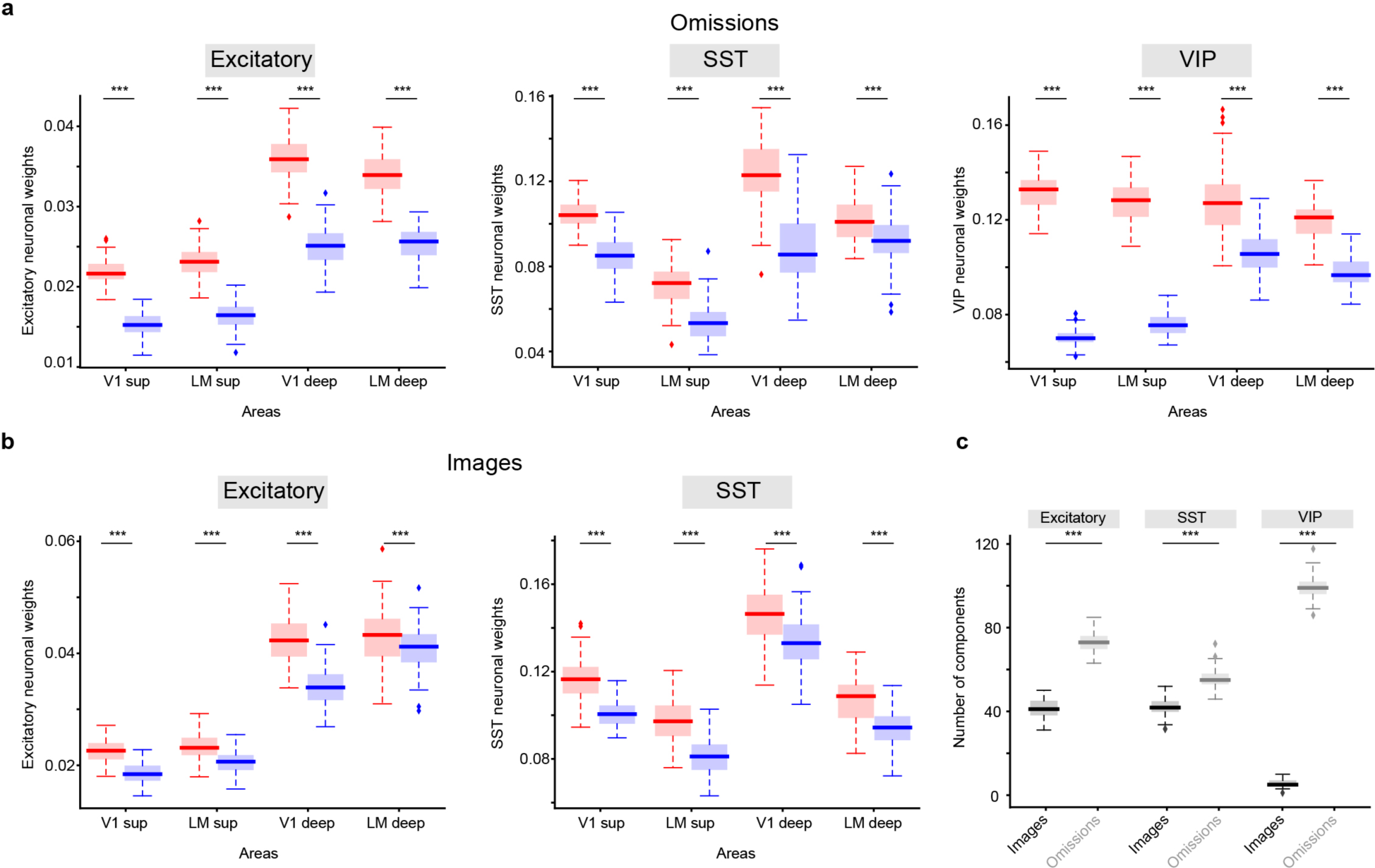
TCA with 10 components. **a, b, and c** depict the results for the TCA analysis for 10 components corresponding to Figure 3D,E,F, respectively.

