## Supplemental materials for "Unexpected events modulate context signaling in VIP and excitatory cells of the visual cortex"

### Supplementary material

| | |
|---|---|
| **Excitatory, V1 image responses** | one-way ANOVA: F= 16.3, p<0.001; TUKEY HSD: p<0.01 for all pairwise comparisons except layer 1-2/3 and layer 1-4. |
| **Excitatory, LM image responses** | TUKEY HSD: p>0.05 for all pairwise comparisons, except for layer 2/3-5: p=0.001. |
| **SST, V1 image responses** | one-way ANOVA: F=12.5, p<0.001; TUKEY HSD: p<0.05 for layer1-2/3, layer 1-4, and layer 4-5 pairwise comparisons. |
| **SST, LM image responses** | one-way ANOVA: F= 1.9, p=0.14. |
| **VIP, V1 image responses** | one-way ANOVA: F=3.0, p=0.03; yet TUKEY HSD: p>0.05 for all pairwise comparisons. |
| **VIP, LM image responses** | one-way ANOVA: F=2.4, p=0.07. VIP, V1 omission responses: one-way ANOVA: F=3.0, p=0.03; yet TUKEY HSD: p>0.05 for all pairwise comparisons. VIP, LM omission responses: one-way ANOVA: F=1.6, p=0.19. |

**Supplementary Table 1.** Statistical tests corresponding to Fig. 1.

| SLC - images | | |
|---|---|---|
| ANOVA | | |
| statistic=974.9242425360012, pvalue=2.053417877366087e-182 | | |
| Posthoc | | |
| Group1 | Group2 | p |
| 0 | 1 | <0.001 |
| 0 | 2 | 0.0045 |
| 0 | 3 | <0.001 |
| 1 | 2 | <0.001 |
| 1 | 3 | 0.17 |
| 2 | 3 | <0.001 |
| Ranksum | | |
| Group | p | effect size |
| 1 | 0.5 | -0.000559821 |
| 2 | <0.001 | 0.005208667 |
| 3 | 0.8 | -0.000677232 |
| 4 | <0.001 | -0.006407717 |

| SLC - omissions | | |
|---|---|---|
| ANOVA | | |
| statistic=924.6397775278477, pvalue=2.0365188317079515e-178 | | |
| Posthoc | | |
| Group1 | Group2 | p |
| 0 | 1 | <0.001 |
| 0 | 2 | 0.9 |
| 0 | 3 | <0.001 |
| 1 | 2 | <0.001 |
| 1 | 3 | 0.9 |
| 2 | 3 | <0.001 |
| Ranksum | | |
| Group | p | effect size |
| 1 | <0.001 | -0.007043246 |
| 2 | <0.001 | -0.010607699 |
| 3 | <0.001 | -0.005858144 |
| 4 | <0.001 | -0.011777935 |

| SST - images | | |
|---|---|---|
| ANOVA | | |
| statistic=562.5678946813928, pvalue=2.34542498531712e-142 | | |
| Posthoc | | |
| Group1 | | |
| 0 | 1 | 0.26 |
| 0 | 2 | <0.001 |
| 0 | 3 | <0.001 |
| 1 | 2 | <0.001 |
| 1 | 3 | <0.001 |
| 2 | 3 | <0.001 |
| Ranksum | | |
| Group | p | effect size |
| 1 | <0.001 | -0.020148597 |
| 2 | <0.001 | 0.008925503 |
| 3 | <0.001 | -0.018708875 |
| 4 | <0.001 | -0.015717922 |

| SST - omissions | | |
|---|---|---|
| ANOVA | | |
| statistic=388.69757476891175, pvalue=1.3433542133282506e-117 | | |
| Posthoc | | |
| Group1 | | |
| 0 | 1 | <0.001 |
| 0 | 2 | <0.001 |
| 0 | 3 | <0.001 |
| 1 | 2 | <0.001 |
| 1 | 3 | <0.001 |
| 2 | 3 | <0.001 |
| Ranksum | | |
| Group | | |
| 1 | <0.001 | -0.021273879 |
| 2 | <0.001 | -0.032506687 |
| 3 | <0.001 | -0.010707941 |
| 4 | <0.001 | -0.061329306 |

| LEGEND | |
|---|---|
| group 1 | LM superficial |
| group 2 | LM deep |
| group 3 | V1 superficial |
| group 4 | V1 deep |

| VIP - omissions | | |
|---|---|---|
| ANOVA | | |
| statistic=152.09786442559385, pvalue=1.4251507357988139e-65 | | |
| Posthoc | | |
| Group1 | | |
| 0 | 1 | <0.001 |
| 0 | 2 | <0.001 |
| 0 | 3 | <0.001 |
| 1 | 2 | <0.001 |
| 1 | 3 | 0.0058 |
| 2 | 3 | <0.001 |
| Ranksum | | |
| Group | | |
| 1 | <0.001 | -0.054864618 |
| 2 | <0.001 | -0.017996513 |
| 3 | <0.001 | -0.068608776 |
| 4 | 0.009 | -0.005409973 |

**Supplementary Table 2.** Statistical tables for TCA analysis.

| Number of components | Celltype | Data | Error | Similarity |
|---|---|---|---|---|
| 5 | Excitatory | Original | 0.9401 | 0.8879 |
| | | Shuffled | 0.9681 | 0.8673 |
| | SST | Original | 0.7695 | 0.9227 |
| | | Shuffled | 0.8365 | 0.9067 |
| | VIP | Original | 0.774 | 0.918 |
| | | Shuffled | 0.8838 | 0.8886 |
| 10 | Excitatory | Original | 0.9084 | 0.8265 |
| | | Shuffled | 0.9366 | 0.8038 |
| | SST | Original | 0.67 | 0.8695 |
| | | Shuffled | 0.7481 | 0.858 |
| | VIP | Original | 0.6991 | 0.8515 |
| | | Shuffled | 0.8238 | 0.8195 |

**Supplementary Table 3.** TCA error and similarity values across with across number of components, celltypes, and original vs shuffled data.

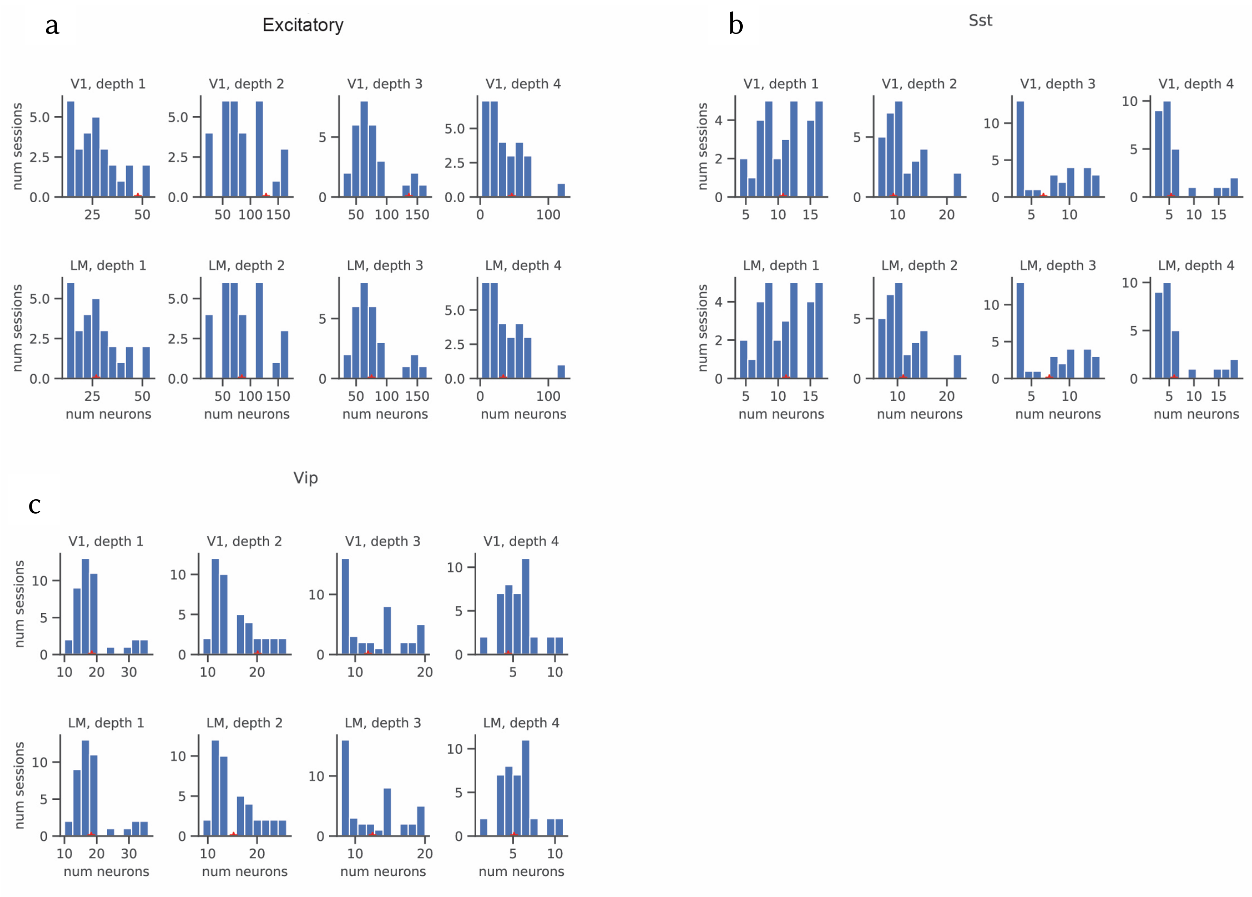


**Supplementary Figure 1. Number of neurons per depth and area, for each cell type.** Distribution of number of neurons recorded simultaneously across depths of V1 (top) and LM (bottom), in all recording sessions, for excitatory **(a)**, SST **(b)**, and VIP **(c)** mice. Note that our goal in this study was to chronically record the activity of distinct excitatory and inhibitory cell types, from the same field of view and across multiple sessions. Like in other studies involving the original Mesoscope[1,2], our system can record from a much larger number of neurons, however, our biological experiments were not designed for that purpose.

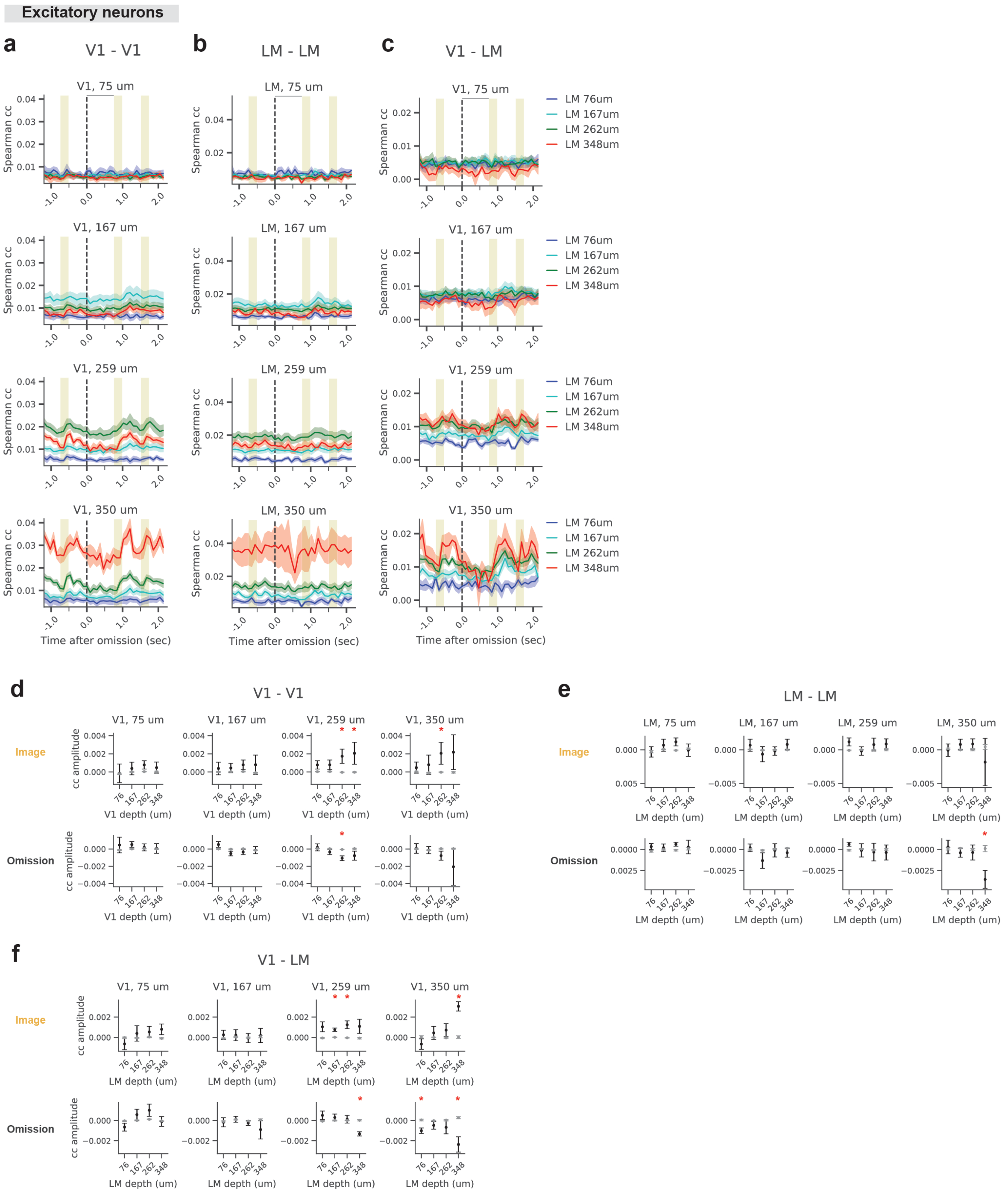


**Supplementary Figure 2. Correlation of excitatory neurons within V1, within LM, and across V1-LM, during images and omissions. a-c** Spearman correlation coefficients computed, at different moments in the trial, between activities of excitatory neurons located in different depths of V1-V1 (a), LM-LM (b), and V1-LM (c). **d-f** Change in correlation coefficients during images (top) and omissions (bottom) relative to the baseline correlation coefficient, computed for the real data (black), and trial shuffled data (gray). Correlation coefficients were quantified over 500 ms after images, and 750 ms after omissions. Red stars indicate statistical significance (two-sided t-test, real vs. shuffle data; $p<0.05$). Traces and error-bars: mean +/- SEM; n = 8 mice.

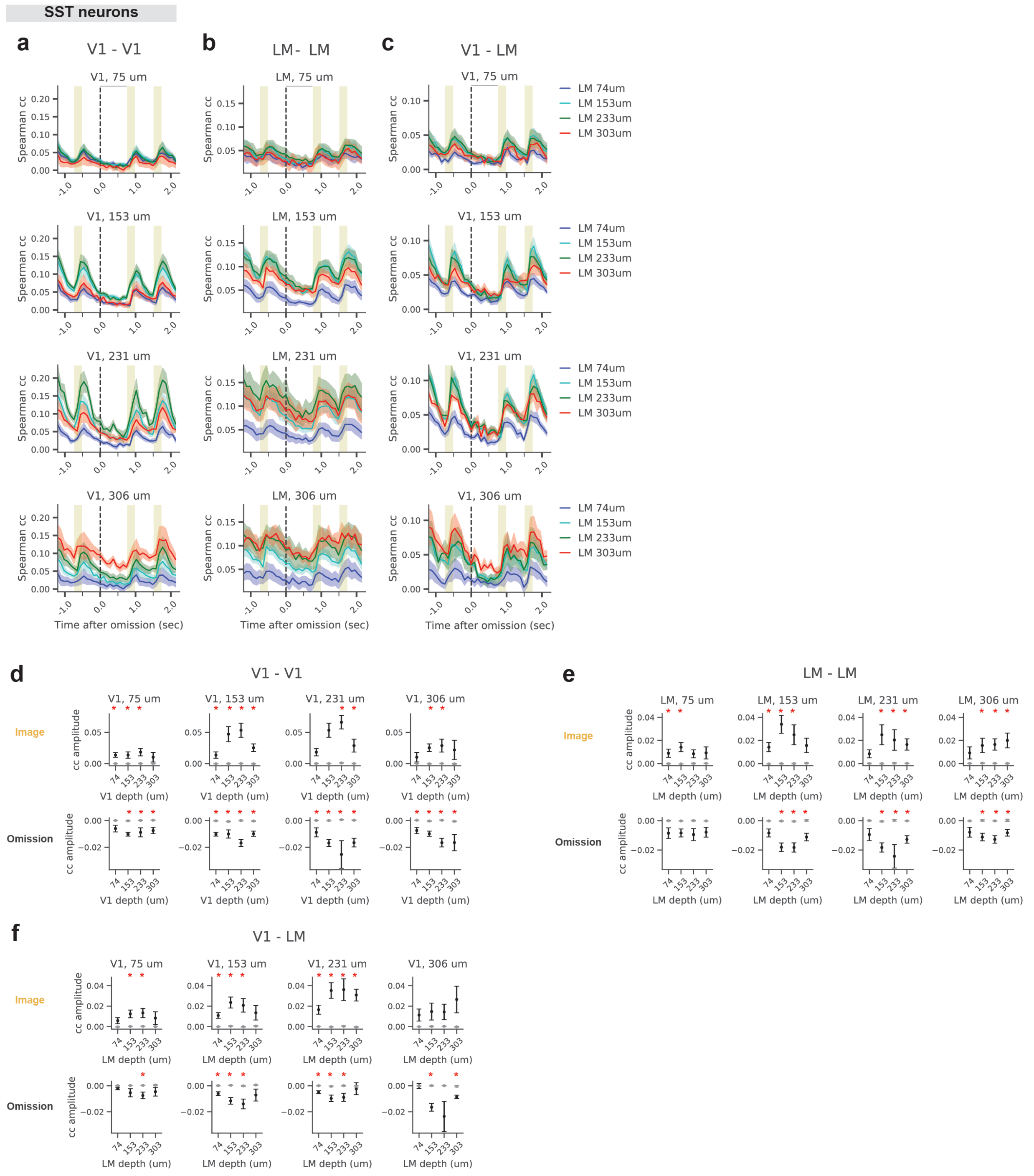


**Supplementary Figure 3.** Correlation **of SST neurons within V1, within LM, and across V1-LM, during images and omissions. a-c** Spearman correlation coefficients computed, at different moments in the trial, between activities of excitatory neurons located in different depths of V1-V1 (a), LM-LM (b), and V1-LM (c). **d-f** Change in correlation coefficients during images (top) and omissions (bottom) relative to the baseline correlation coefficient, computed for the real data (black), and trial shuffled data (gray). Correlation coefficients were quantified over 500 ms after images, and 750 ms after omissions. Red stars indicate statistical significance (two-sided t-test, real vs. shuffle data; p<0.05). Traces and error-bars: mean +/- SEM; n = 6 mice.

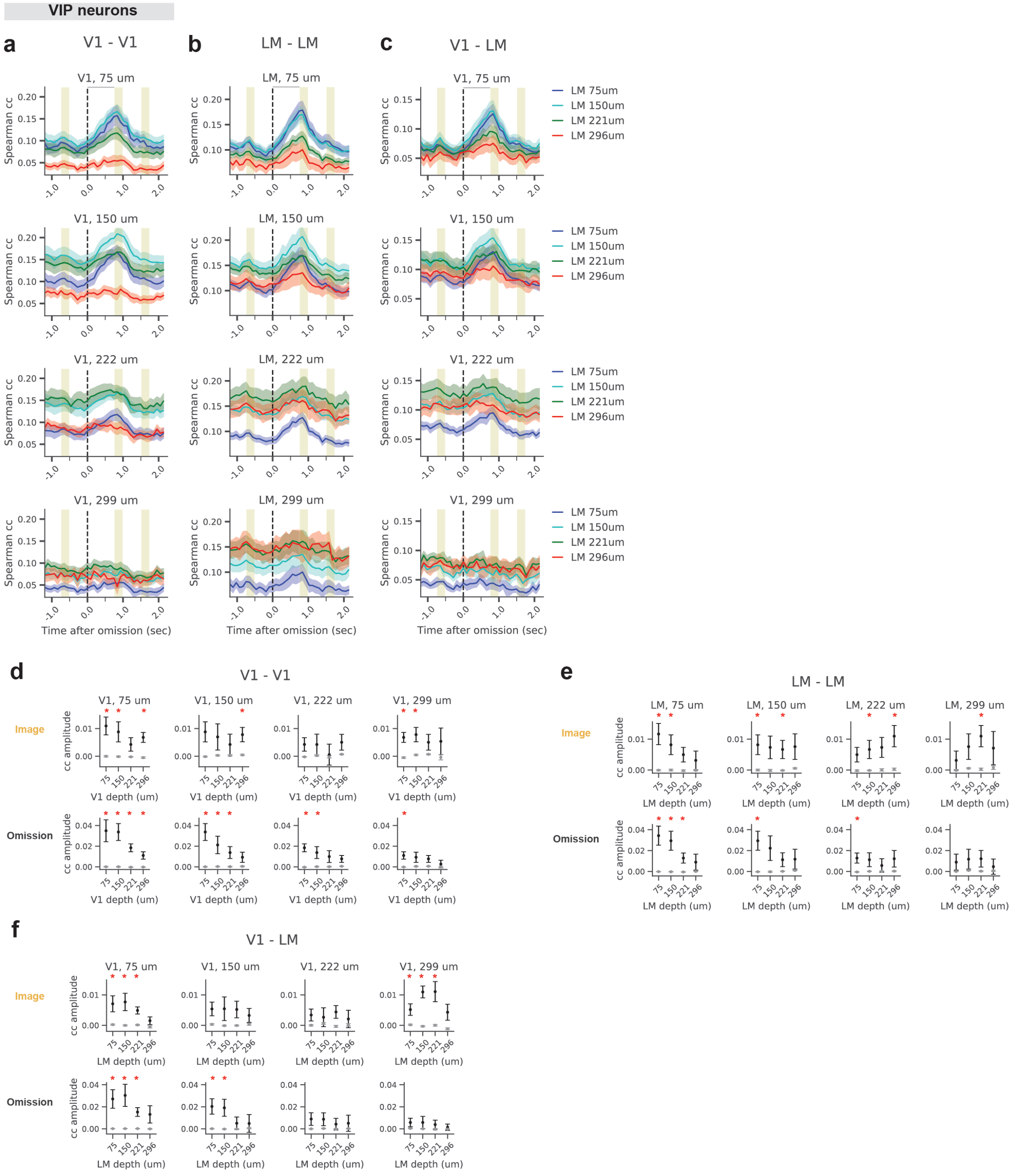


**Supplementary Figure 4. Correlation of VIP neurons within V1, within LM, and across V1-LM, during images and omissions. a-c** Spearman correlation coefficients computed, at different moments in the trial, between activities of excitatory neurons located in different depths of V1-V1 (a), LM-LM (b), and V1-LM (c). **d-f** Change in correlation coefficients during images (top) and omissions (bottom) relative to the baseline correlation coefficient, computed for the real data (black), and trial shuffled data (gray). Correlation coefficients were quantified over 500 ms after images, and 750 ms after omissions. Red stars indicate statistical significance (two-sided t-test, real vs. shuffle data; $p<0.05$). Traces and error-bars: mean +/- SEM; n = 9 mice.

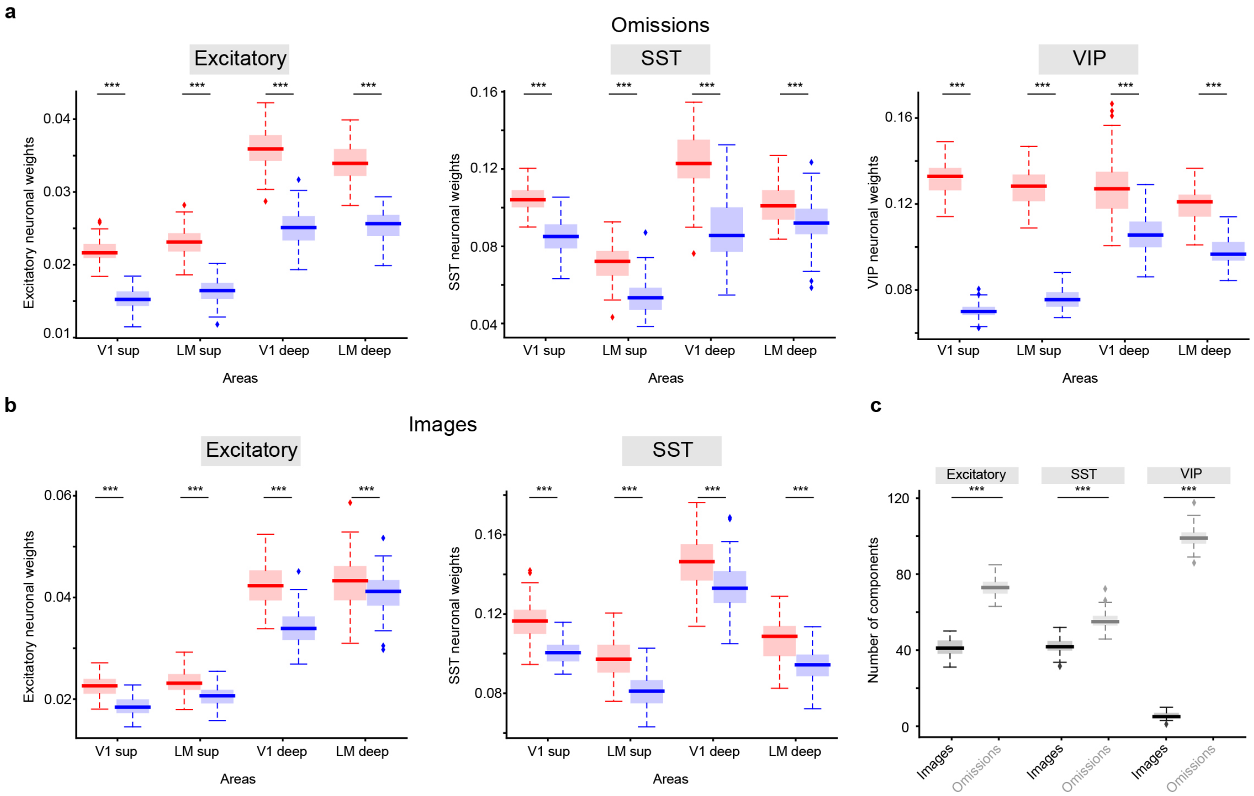


**Supplementary Figure 5. TCA with 10 components. a, b, and c** depict the results for the TCA analysis for 10 components corresponding to Figure 3D,E,F, respectively.
